# Multiple molecular links between the circadian clock and memory centers in honey bees

**DOI:** 10.1101/2024.03.31.587450

**Authors:** Tiyasa Roy, Rikesh Jain, Axel Brockmann

## Abstract

Time and memory are intimately linked: the capability to learn and recall varies over the day and humans and many animals can associate important events with the time of day. However, how the circadian clock and memory centers are connected is not well understood. We time-trained honey bee foragers and used RNA-sequencing and RNAscope imaging to analyze gene expression changes in focal populations of mushroom body neurons. Thus, we identified three candidate functional modules of time-memory: synchronized peak-level expression of memory-related genes during training time, anticipatory activation of transcription in *pdfr*-expressing neurons, and *cry2* and *per* co-expressing neurons that might represent local clocks. The complex interactions between the clock and memory centers, which appear to be more similar to mammals than other insects, might have been facilitated to optimize social foraging in honey bees.

## Introduction

Many animals can memorize the time-of-day they made a particular experience (*1-4*), which allows them to anticipate recurrent events and prepare appropriate behavioral responses in advance (*5,6*). Learning and recalling the time-of-day require a functional connection between the circadian clock and brain regions involved in learning and memory (*6,7*). Previous studies in *Drosophila* and mice have focused on daily oscillation of gene expression coinciding with changes in the capability to form memories and genetic manipulation experiments demonstrating an effect of clock genes on time-of-day memories (*4,8-10*). However, how signals from the clock and sensory systems are combined and consolidated in memory cells is still unclear. Here, we used social honey bees which have elaborated mushroom bodies (MBs), the major insect memory centers (*11*), and show an exceptional capability to form multiple time-memories under free-flight conditions (*12,13*) to search for candidate molecular interactions between the clock and memory centers. Our project was motivated by an earlier study from our lab showing that expression of the neuronal-activity regulated transcription factor (TF) *Egr1* in the small Kenyon cells (sKCs), the major multimodal neuronal population of the MBs, correlates with training-time (*14,15*). Using laser-capture microdissection (LCM) of sKC-enriched tissue samples and RNAscope imaging (an advanced type of single molecule fluorescent in situ hybridization, smFISH) we were able to detect for the first-time spatiotemporal expression dynamics of multiple genes in a selected subset of MB neurons that allowed us to address the following questions: (i) Do genes in the sKCs show expression changes associated with the training-time? (ii) Do sKCs receive input from the central pacemaker (CP), the master circadian clock, and (iii) do KCs express clock genes that would enable them to function as local clocks?

## Results

### Daily gene expression dynamics in sKCs of time-trained honey bee foragers

Foragers of a honey bee colony allocate their daily foraging activities according to peak times of food availability (*16*). Experimental feeder time-training synchronizes the behavioral activity and physiological processes among foragers visiting the same feeder (*17*). Thus, time-training not only initiates molecular processes involved in time-memory but also synchronizes temporal dynamics of gene expression among individuals (*14,18,19*). We trained honey bee (*Apis mellifera*) foragers to visit a feeder from 09:00 to 11:00 (1M sucrose solution) in an outdoor flight enclosure for 10 consecutive days (Fig. 1A and methods). Foragers visiting the feeder on at least 3 out of the last 4 training days (indicated by different color marks for each day) were considered as time-trained and then we collected them in liquid nitrogen on day 11 at 7 different time-points [06:00, 10:00, 14:00. 18:00, 22:00, 02:00, 06:00, (Fig. 1B)], while the feeder was presented but not rewarded, omitting reward induced gene expression changes (Fig. 1, A and B, also see methods).

**Fig. 1.**
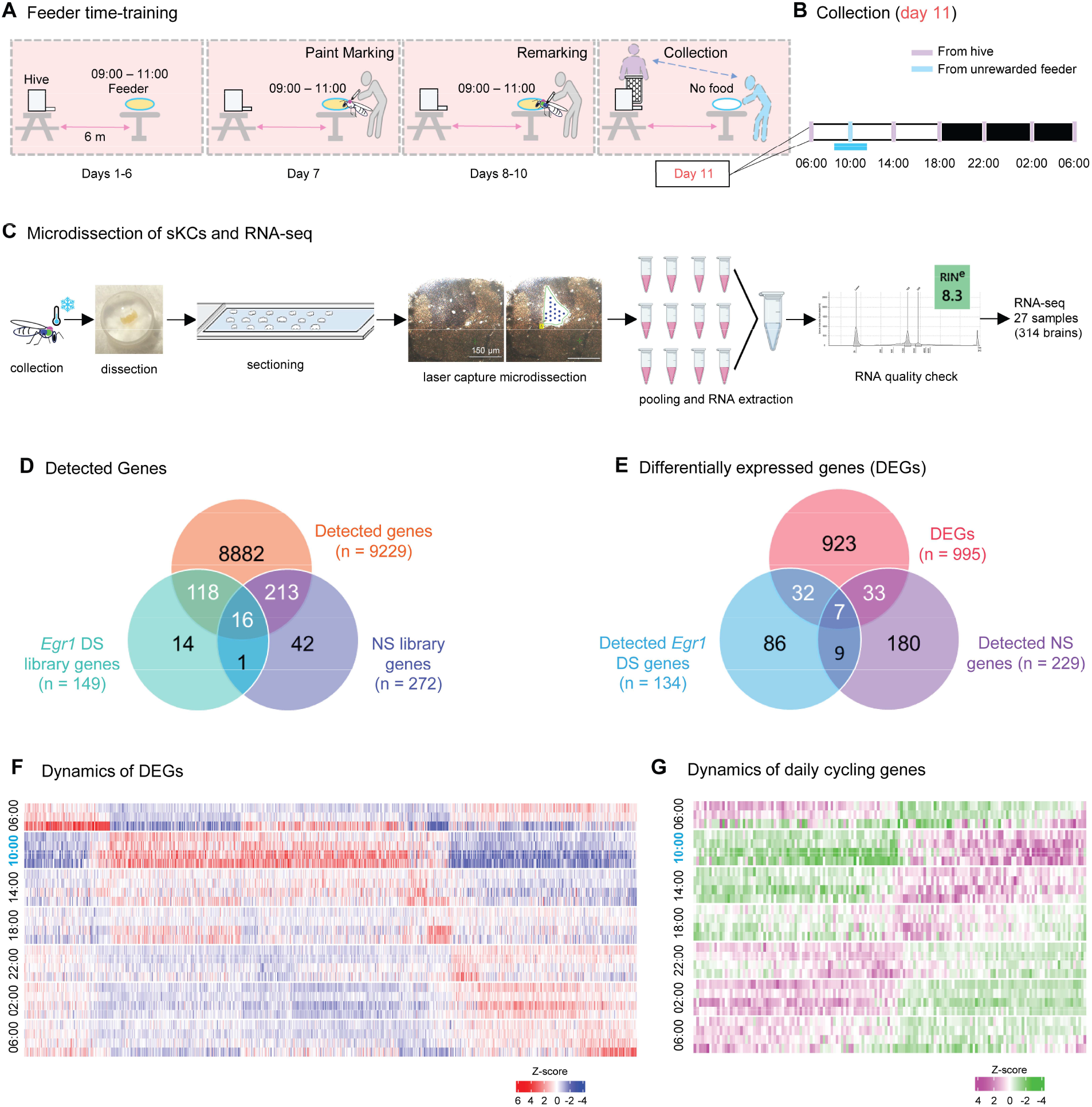
SKC-enriched transcriptome of time-trained foragers. (**A** and **B**) Schematic illustration of feeder time-training paradigm (A) and collection time-points (B). (**C**) Overview of experimental pipeline for RNA-seq with sKC-enriched population. For (A) to (C) see methods. (**D**) Venn diagram showing the overlap among total number (no.) of detected genes, libraries prepared for NS genes (literature search) and *Egr1* DS genes (bioinformatically predicted and provided by authors;*28*). (**E**) Venn diagram depicting overlap among triple sets, i.e., total no. of DEGs, detected NS- and *Egr1* DS-library genes. (**F**) Heatmap showing temporal expression pattern of all the DEGs. (**G**) Heatmap for expression dynamics of 152 genes showing circadian oscillation (daily periodicity) in MetaCycle algorithm, Fisher’s exact test, BH.Q < 0.05. See also fig. S4B. For all heatmaps consecutive two columns for each collection time point represent replicates from same colony, and next two columns from a second colony. Both colonies are independent and the behavioral experiments were performed separately.

For identifying genes involved in time-memory we collected sKC-enriched tissue samples (*14*). The sKC cell bodies are closely arranged at the inner core of the MB calyces, and this area is visually distinguishable from that of the lKCs (*20*). We dissected the sKCs of all four MB calyces of one brain using LCM and pooled the RNA samples from 11-12 brains to yield high-quality RNA. For each of the 7 different collection times, we collected two replicates each from two different colonies for RNA-sequencing [RNA-seq (Fig. 1C), see methods]. Finally, we sequenced a total of 27 sKC-enriched RNA samples as we lost one sample due to technical issues. We consistently detected reads for 9229 genes (filtering criteria = minimum count value of 5 in one or more replicates per time-point, data S1). Gene ontology (GO) analysis of all detected genes (DGs) revealed enrichment of TFs, ribosome and initiation factors, SNARE interactions in vesicular transport and ER-Golgi transport and protein kinases, among others [FDR < 0.01; (fig. S1, A and B)]. To refine our search for candidate time-memory genes we prepared two gene libraries based on published data (see data S2): (i) *Egr1* and its candidate downstream genes (*Egr1* DS, n=149) and (ii) neuronal signaling genes (NS, n=272) including neurotransmitters, receptors, second messengers etc. Among the total number of the 9229 DGs, we found 134 out of 149 *Egr1* DS genes (90%), and 229 out of 272 NS genes (84%) (Fig. 1D and fig. S2, A to I).

Next, we asked how many of the 9229 DGs are differently regulated and how many of those show an expression pattern aligned to the training-time or a circadian oscillation. First, we performed a differential gene expression analysis (pairwise comparison test, p.adj < 0.05, for details see methods) and identified 995 genes (11% of all detected genes) to be differentially expressed (Fig. 1, E and F, data S3). A GO analysis for these differentially expressed genes (DEGs) showed an enrichment of ecdysteroid kinase, motor protein domains, G-protein coupled receptor gene sets among others [FDR < 0.01; (fig. S3, A and B)]. Arranging the 995 DEGs according to peak expression values (Fig. 1F) revealed that 862 (86.6% of all DEGs) genes show maximum (n = 484) or minimum (n = 378) peak expression at 10:00, the collection time-point which is one hour into the feeder training time (Fig. 1F and fig. S4A). Finally, we used the MetaCycle algorithm to examine which of the 9229 DGs might exhibit circadian oscillations in gene expression. We found only 152 (1.6%) genes showed a significant daily oscillation at BH.Q<0.05 (Fig. 1G); and lowering the threshold to BH.Q<0.3 increased the number to 686 (7.4%) genes (fig. S4B) with Fisher’s exact test.

To summarize, our analyses indicate that the temporal expression dynamics in the sKCs of time-trained honey bee foragers more likely follow the training schedule than a general daily circadian rhythm (Fig. 1, F and G and fig. S4, C and D). Particularly, the high ratio of DEGs showing peak expression during the training time suggest that the expression pattern is a result of the time-training, likely regulated by *Egr1* in combination with other TFs (*14*).

### Candidate genes involved in molecular processes underlying time-memory

Similar to the majority of all DEGs, most of the *Egr1* DS (35 out of 39, 89.7%) and NS (31 out of 40, 77.5%) DEGs exhibited peak expression at 10:00 (training-time) (Fig. 2, A and B). Among the 35 *Egr1* DS DEGs with highest expression at 10:00, 5 genes are involved in dopamine/ecdysteroid signaling (e.g. *DopEcR, Ddc, E74*), 10 genes in intracellular signaling cascades (e.g. *cac, meng-po*), 5 genes in regulating or modulating transcription (e.g. *Atf3, Spt6, Imp*), and 3 synaptic genes (e.g. *snap25, tutl*). In addition, 3 of the genes peaking at 10:00 (*Ap-1, SIK2, Inos*) have been reported to function in modulating the core molecular clock in both insects and vertebrates (*21-23*). In the group of 31 NS DEGs with peak expression at 10:00, we found 12 genes functioning in neurotransmitter and biogenic amine signaling (e.g. *nAChRa, mGlutR1, TyrR, Octbeta2, Dop1*), 6 genes involved in neuropeptide signaling (e.g. *AstC, SIFR*), 6 genes of intracellular signaling cascades (e.g. *cac, Mpk3*), 6 TFs (e.g. *AP-1, Egr1, Hr38, Creb*), and 2 synaptic genes (*Snap25, Syx1A*). Several of the NS DEGs have previously been reported to play vital roles in learning and memory processes in honey bees and other organisms [e.g. octopamine and glutamate signaling, cAMP-PKA cascade and *Creb* (*24-26*)].

**Fig. 2.**
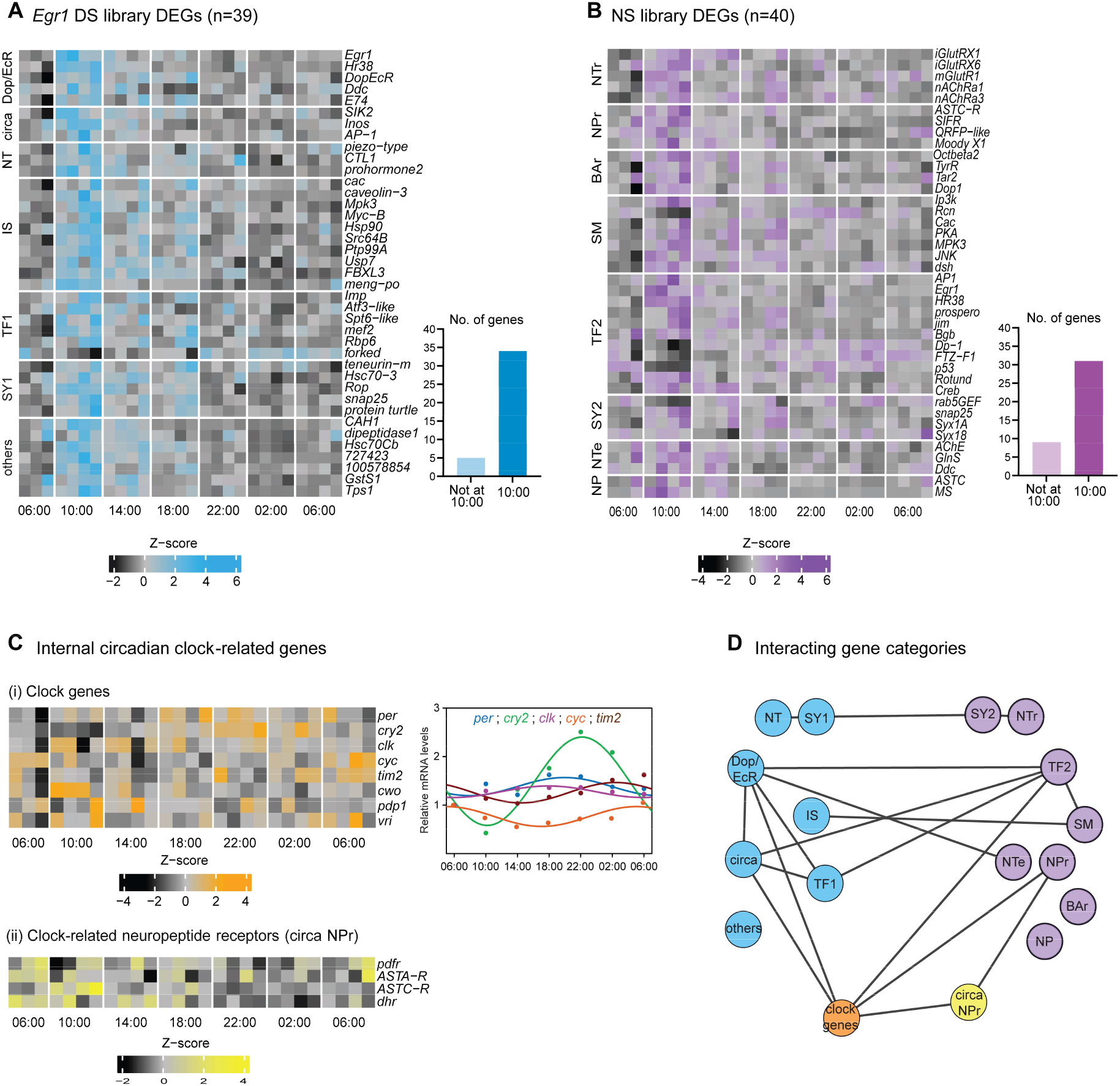
Correlation between training time and gene expression pattern of focal groups of genes. (**A** and **B**) Heatmaps showing expression of *Egr1* DS and NS library DEGs. Corresponding bar diagrams showing 89.74% *Egr1* DS DEGs and 77.5% NS DEGs at their highest expression at 10:00. The genes are separated into different categories based on literature review. In (A): Dop/EcR, dopamine-ecdysteroid signaling; circa, genes involved in molecular circadian clock; NT, neurotransmission; IS, intracellular signaling; TF1, transcription factors; SY1, synaptic genes; others, genes that could not be designated into a functional category. In (B): NTr, receptors of neurotransmitters; NPr, receptors of neuropeptides; BAr, receptors for biogenic amines; SM, second messenger cascade genes; TF2, transcription factors; SY2, synaptic genes; NTe, neurotransmitter enzymes; NP, neuropeptide genes. (**C**) (i) Heatmap showing expression of all known canonical clock genes. Right panel: Cosinor plot showing expression of 5 clock genes characterized for whole brain in (*31*). (ii) Heatmap showing expression of clock-related neuropeptide receptors. (**D**) Illustration of gene networks showing interaction between any two categories if one or more genes overlap for *Egr1* DS- and NS-DEGs and internal circadian clock-related genes.

In addition, we screened the total DGs for two groups of genes which are highly likely involved in time-memory (i) genes of the ecdysteroid signaling pathway and (ii) circadian clock genes. Altogether we found 9 genes of the ecdysteroid signaling pathway, 4 of them (*DopEcR, E74, Egr1, Hr38*) showed changes in gene expression according to training time and 5 genes (*EcR, Usp, E75, Br-c, Mblk1/E93*) did not show any change in expression (fig. S2A). For all these genes, except *DopEcR*, expression in sKCs or lKCs has been reported before (*27*) and several studies in honey bees have documented an important role of ecdysteroid-signaling in the onset of foraging, active foraging, and learning and memory during foraging (*14,18,28,29*). Also, evidence in *Drosophila*, indicates that ecdysteroid-signaling (e.g., *E75*) is involved in modulating the core molecular clock (*30*).

Further we found all the core clock genes (*cry2, per, clk, cyc, tim2, cwo, pdp1* and *vri*) among the DGs expressed in the sKC-enriched tissue samples. Only *cry2* exhibited a significant expression difference (p.adj < 0.05, peak expression at 22:00, Fig. 1F, 2C) and also significant circadian oscillation (MetaCycle, BH.Q< 0.05, Fig 1G, 2C). Both findings are consistent with whole brain mRNA expression data for honey bees trained to forage at a feeder in the morning (*31*). Further, we found *pdfr*, the receptor that binds pigment-dispersing factor (PDF), which is regarded as the major output neuropeptide exclusively produced by the central pacemaker (CP) neurons in many insects (*32-34*) as well as 9 other neuropeptides and neuropeptide receptors (*AstA, AstC, Dh31, Dh44*) which have been reported to connect different clock neurons as well as clock neurons and target cells in *Drosophila* [(*35*), Fig. 2C and fig. S5A)].

A first step gene network analysis (Fig. 2D) based on the gene categories used for our analyses suggests that interactions between clock genes, genes of the dopamine/ecdysteroid signaling pathway and multiple TFs prime the molecular processes generating time-memories in honey bees.

### Co-expression of Egr1 and pdfr in MBs indicates direct input from the CP

To explore whether there is a functional connection between *Egr1* expressing KCs and the circadian clock, we concentrated on two questions: (i) do sKCs show *Egr1* expression in anticipation of the training-time indicating that the expression is regulated by an internal mechanism and (ii) do those sKCs that express *Egr1* co-express *pdfr* confirming a direct input from the CP. The idea for this experiment was based on two earlier findings. First, behavioral studies on time-memory in honey bees showed that time-trained foragers exhibit a characteristic anticipatory activity involving moving towards the entrance area of the hive and early flights to the feeder (*19,36*). Second, a molecular study in our lab revealed that *Egr1* expression is initiated in sKCs during the time of the training phase even if the foragers are prevented from leaving the hive indicating that the initiation of the expression is independent of foraging-related cues (*14*). For the study of *Egr1* and *pdfr* expression we performed RNAscope imaging using custom-made probes (see methods). The feeder time-training was carried out as in the RNA-seq experiment, except that on the collection day the feeder contained the food reward to describe the spatiotemporal expression changes from anticipation to continuous foraging and we collected foragers at three different time-points: 08:00 (anticipatory phase, i.e., when the foragers were still inside the colony but had moved to the entrance, N=3) 09:00 (start of feeder time-training and foraging, N=3), and 10:00 (continuous foraging, N=3).

At 08:00, *Egr1* and *pdfr* expression was already initiated but almost exclusively restricted to the sKCs region (Fig. 3A). At 09:00, *Egr1* and *pdfr* expression extended all over the sKC population, some parts of the mKCs, and only small parts of the lKCs (Fig. 3B) and by 10:00 *Egr1* and *pdfr* expression was activated over the whole cell body area of the MB calyces (Fig. 3C). mRNAs of *Egr1* and *pdfr* were detected in- and out-side of DAPI staining indicating that both transcripts are transported outside the nucleus likely for the synthesis of the respective proteins [(Fig. 3D), and for specificity and efficiency of RNAscope see fig. S6, A to C]. For all the three times-points, we detected significant co-expression of *Egr1* and *pdfr*, measured as high degree of overlap between the fluorescent signals of the two probes (Fig. 3E; mean Pearson’s r ≥ 0.5, for Manders’ M1 and M2∼1.0; fig. S7, A and B). That the area showing co-expression expanded from few KCs to majority of the KCs across the three time-points was confirmed by quantitative analysis (Fig. 3F). The staining pattern indicates that the majority of sKCs, mKCs, and lKCs co-express *Egr1* and *pdfr* and thus receive direct neuromodulatory input from the CP. Further, expression of both genes in some KCs during the anticipatory phase when the bees were still inside the colony suggests that at least in these cells expression of *Egr1* could be solely activated via input from the CP (Fig. 3, F and G).

**Fig. 3.**
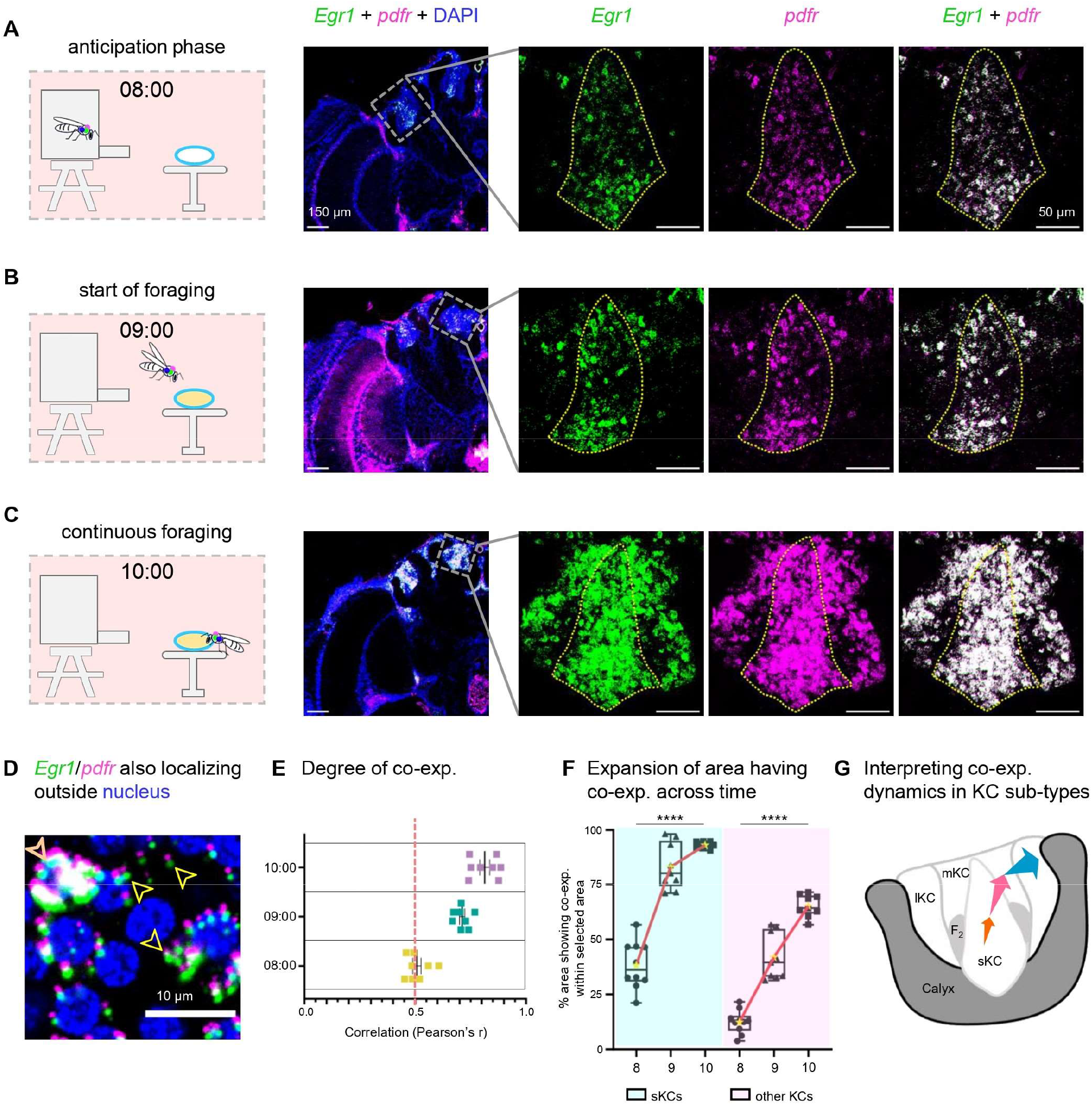
*Egr1* and *pdfr* expression dynamics in MBs of time-trained foragers. (**A**) RNAscope images for expression of *Egr1* (green) and *pdfr* (magenta) preferentially restricted to the sKCs (dashed yellow line) at 08:00, anticipation phase of feeder time-training. (**B**) RNAscope images showing *Egr1* (green) and *pdfr* (magenta) are co-expressed in more KCs compared to (A) i.e., in more sKCs, and beyond sKCs at 09:00, start of active foraging. (**C**) RNAscope images showing co-expression of *Egr1* (green) and *pdfr* (magenta) is expanded to mostly all KCs apart from a specific KC area at 10:00, one hour into active foraging. For (A), (B) and (C) right panel shows magnified image of one calyx (as indicated by marked area of left panel). Scale bars, lower magnification images (left panel, showing one hemisphere of brain), 150 μm. Scale bars, higher magnification images (right panel, showing one calyx), 50 μm. Brain section depth: Mid, 150 to 300 μm. Observations were universal over: No. of individuals (N) = 3, and no. of sections per brain (n) = 15 for each of the three time points. (**D**) Magnified image up to single cell (nuclei) resolution where colors are used for respective genes. *pdfr* (magenta) and *Egr1* (green). Puncta (transcripts) are either localized within the nucleus; top-left: orange arrowhead (blue, DAPI) or outside the nuclear area; yellow arrowheads. (**E**) Scatter dot plot (mean and SEM) showing Pearson’s correlation coefficient (r) for time-points, 08:00, 09:00, and 10:00, suggesting high degree of correlation between *Egr1* and *pdfr* (r ≥ 0.5). (**F**) Box-whiskers plot showing percentage (%) of co-expressed (co-exp.) pixels for *Egr1* and *pdfr* in the sKC area (blue) and other KCs (pink) at 08:00 (8), 09:00 (9) and 10:00 (10). The trendline joins the means (yellow stars) of each column. Test for linear trend in one-way ANOVA, p<0.05 stands significant (****, p<0.0001). For both (E) and (F) different calyces at high magnification were used for quantification across N=3 at equivalent brain depth. The respective KC areas were identified by DAPI staining. (**G**) Schematic representation of KC sub-types (*11*) and highlighted in orange, pink, and blue the expansion of preferential co-exp. from sKCs to more KCs (sKCs + mKCs) to all KCs (sKCs + mKCs + lKCs) except *foxP* ^+^ region (fig. S8) at 08:00, 09:00 and 10:00, respectively color-coded.

### Co-expression of clock genes suggests local clocks in memory centers

As our RNA-seq study detected clock gene expression in the sKC-enriched tissue samples the questions arose which KCs express clock genes, and whether at least some of those co-express the two major clock genes *per* and *cry2*. Co-expression of both genes paired with oscillating RNA expression would be strong evidence that those cells exhibit a functional molecular clock (*37*). Similar to our first smFISH experiment we used custom made RNAscope probes for *per, cry2* and *pdfr*. First, staining with the *per* ISH probes detected 3 clear populations of neuronal cell bodies between the protocerebrum and the optic lobes that were previously identified as the dorsolateral neurons (DLN) and lateral neurons 1 and 2 (LN_1_, LN_2_). These cells are located at a similar ventral-dorsal position in the brain and comprise the majority of the CP (Fig. 4A; *33*). Confirming this identification, we also found co-expression with *cry2* and *pdfr* in these cells (Fig. 4A). In the MB calyces, we found widely distributed low expression levels of *per, cry2* and *pdfr* with local areas showing higher mRNA levels particularly of *per* and *cry2* and a higher incidence of co-expression (see orange arrows in Fig. 4B; left panel and Fig. 4C). These areas occurred in three major cell groups within a calyx (a sKCs group, and in each of the two areas of lKCs adjoining the sKCs, Fig. 4C and fig. S9, A and B). Recent studies with a polyclonal antibody against PER only detected a few cells at the bottom of the calyces (*33*, table S1). The level of *cry2* expression in these clusters was high at 22:00 and low at 10:00 consistent with the oscillation of *cry2* in our RNA-seq data and matching the general whole brain oscillation of *cry2* of morning-trained foragers (fig. S9C; *31*). The location of the clusters in the lKCs appears to be consistent among the brains of different individuals, whereas the location of the sKC cluster shows interindividual variation (Fig. 4B and fig. S9A). The heightened levels of *per* and *cry2* expression and increased visual signs of co-expression of *per, cry2* and *pdfr* in these clusters, resembling the staining of the CP, is strong evidence that some of the memory cells in honey bee MBs function as local clocks (fig. S9, D to F). On the other hand, the wide distribution of low-level expression of the clock genes and interindividual variation in the location of the clusters of heightened expression suggest that these cells do not represent a specific type of KCs but that the majority or at least a large group of KCs have the capability to become clock cells.

**Fig. 4.**
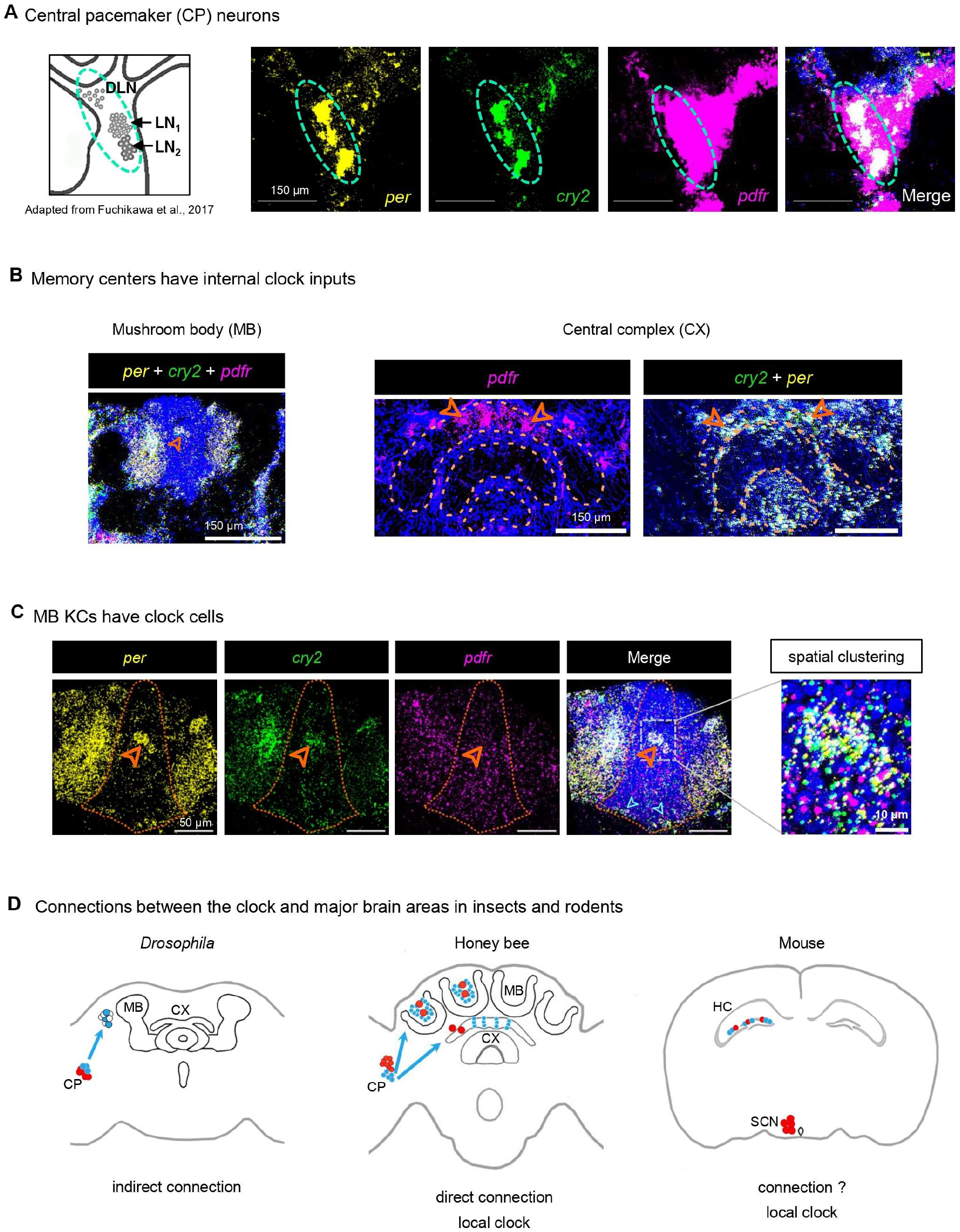
Local circadian clock in MB KCs and central complex. (**A**) RNAscope images showing expression of *per* (yellow), *cry2* (green), and *pdfr* (magenta) and co-expression (white) in CP neurons. DAPI, blue. Left panel: Schematic representation of identified CP neurons in honey bee foragers (*33*). (**B**) RNAscope images showing expression of circadian clock inputs in memory centers. Left major panel: *per* (yellow), *cry2* (green) and *pdfr* (magenta) expressions in MB. Right major panel: *pdfr* expression in central complex (left), and expressions of *per* (yellow) and *cry2* (green) in cell groups in close vicinity of central complex (right). Scale bars, as indicated in respective panels. DAPI, blue. (**C**) RNAscope images for *per* (yellow), *cry2* (green), and *pdfr* (magenta) and co-expression (white). sKC region, orange dotted line. Arrowheads: orange, spatial clustering of cells showing high transcript abundance; cyan, *per* (yellow) and *cry2* (green) co-expression in basal sKCs indicated earlier with immunohistochemistry with PER in (*33*). Right panel: High magnification image showing cells with high transcript abundance spatially clustered. DAPI, blue. (**D**) Schematic representation showing current state of understanding of functional interaction between circadian clock and memory centers in *Drosophila* (earlier studies; *45*), in honey bees (this study) and rodents (earlier studies; *43,47*). SCN, suprachiasmatic nucleus; HC, hippocampus.

Apart from the MBs, we also found *pdfr, per*, and *cry2* expression in cell groups associated with the central complex, the other higher order brain neuropil in insects, involved in e.g. navigation, time-compensated sun compass orientation and dance communication (*38-40*). Expression of *pdfr* could be clearly detected in groups of cell bodies posterior to the protocerebral bridge (PB) that are arranged above each other similar to the cell bodies of the columnar neurons that are part of the sun-compass system (*41*, Fig. 4B; right panels). Co-expression of *cry2* and *per* occurred in cell groups in close vicinity of the PB and central body (Fig. 4C), but whether they belong to the central complex requires more detailed studies.

## Discussion

Combining time-training of honey bee foragers under natural conditions with time-series RNA-seq and RNAscope imaging of a defined subset of MB neurons we identified three basic functional modules of time-memory: (i) anticipatory activation of gene transcription in sKCs that express a receptor for the major output signal of the circadian clock, (ii) synchronized peak-level expression of memory-related genes during the time of training, and (iii) memory cells co-expressing *per* and *cry2* that enables them to function as local clocks.

Time-trained honey bees show anticipatory activity involving aggregating near the hive entrance and early flights toward the food source (*36*). Stimulation with a familiar odor during this phase was shown to activate navigational memory (*42*). Thus, anticipatory activation of sKCs by CP, that also regulates foraging activity, could function in a timely reactivation of navigational memory and preparation for memory reconsolidation. Previous studies in flies and mice already demonstrated an important role of major clock genes in the timing of memory processes (*3,4,9*). Widespread co-expression of *pdfr* and the TF *Egr1* in KCs indicates a second, neuromodulator based, connection between the clock and memory center (*14*). As *Egr1* expression in honey bees correlates with time-training schedules, *Egr1* activation via Pdf might play an important role in their ability to form multiple time-memories. A neuromodulator connection between the SCN and hippocampus has been suggested but not yet demonstrated and in *Drosophila*, it appears that the CP is only indirectly connected to the MBs (*8,43-46*) (Fig. 4D). Finally, memory cells functioning as possible local clocks have first been described in the hippocampus of mice and it was hypothesized that they could serve two functions: autonomous timekeeping supporting regulation of gene expression independent of the CP and general daily activity patterns (*46,47*) or providing time-of-day information that could be combined with other contextual features of an event like the color or odor of a flower (*7*). Our study reports for the first-time anatomical evidence for local clocks in insect MBs.

The diverse molecular interactions between the circadian clock and MBs in honey bees corresponds with the varied behavioral experiments indicating a highly elaborated capability to form multiple time-memories per day (*12*; Fig. 4D). This capability was associated to differences in flowering times but there is increasing evidence that it evolved to optimize spatiotemporal foraging activities in social insect colonies (*14,16,48*). This exceptional natural behavior makes honey bees a most convenient animal model to explore fundamental principles and mechanisms of time-memory (*7,49,50*).

## Acknowledgements

We thank Charlotte Helfrich-Förster, Wulfila Gronenberg, and Gene E. Robinson for their comments on the manuscript. We also thank the Next Generation Genomics Facility at NCBS, particularly Awadhesh Pandit for assisting with the RNA-seq run. We are grateful to the Central Imaging and Flow Cytometry Facility (CIFF) and the instrumentation team at NCBS for their support. Further, we acknowledge the technical advice and support from Zeiss Microscopy team and Advanced Cell Diagnostics (ACD). We are thankful to Lukumoni Das, Sukrithi N Venu and Sripriya Bulusu for helping with the behavioral experiments.

## Author Contributions

Conceptualization: T.R., A.B.; Methodology: T.R., A.B.; Investigation: T.R.; Visualization: T.R., R.J., A.B.; Formal analysis: T.R., R.J., A.B.; Funding acquisition: A.B.; Supervision: A.B.; Writing-original draft: T.R., A.B.; Writing-review and editing: T.R., A.B.

## Data availability

The RNA-seq data will be published in NCBI Sequence Read Archive upon acceptance for publication.

## Materials and Methods

### Animals

*Apis mellifera* colonies consisting of a naturally mated queen and around 8000 workers (i.e. 8 frames with ∼1000 workers per frame) were procured from a local beekeeper and maintained on the campus of National Centre for Biological Sciences-Tata Institute of Fundamental Research (NCBS-TIFR), Bangalore, India. For all experiments colonies were kept in an outdoor flight cage (12m x 4m x 4m), which allowed to control time of food availability while otherwise exposing the bees to the natural light/dark cycle. The experiments were performed in September and October 2021, when sunrise and sunset started around 06:10h and 18:05h respectively (timeanddate.com). Thus, the bees experienced an approximate 12:12 LD cycle.

### Time-training paradigm

Colonies were allowed to adjust to the flight cage for 2-3 days prior to the start of the experiment. During this pre-training period, 1M sucrose solution and pollen were provided ad libitum on two unscented but differently colored feeders (09:00 to 17:00). Then during the time-training only the sucrose feeder (1M sucrose solution) was presented for 2 hours from 09:00 to 11:00. Foragers were trained for 10 consecutive days to improve temporal accuracy of feeder visits (*51*) and increase the total number of foragers, as a huge number of individuals were needed for the molecular study. Each day at the end of the training period the feeder plate was cleaned and put back at its location to prevent association of feeder plate with the presentation time for food reward (*18*). On the 7th day all foragers visiting the feeder, were marked at the feeder during the training period. From day 8 to day 10, only those already marked foragers were remarked with different colors on each day when visiting the feeder plate during the 2 hours training period. On the collection day, we only caught consistent foragers (*51*) that had visited the feeder on three of the four previous days, i.e. showing at least three different color marks. Further, we chose a training time period from 09:00 to 11:00 because the anticipatory activity phase of time-trained foragers is shorter in the morning (*36*) which might further help to increase the level of physiological synchronization among the experimental group of foragers.

### Collection of time-trained foragers

On the 11th day, food was not provided at the feeder plate. Marked consistent foragers were collected at 7 different time-points at regular 4h intervals i.e., at 06:00, 10:00, 14:00, 18:00, 22:00, 02:00, and 06:00 for a complete circadian day (24h) starting at 06:00 (11th day) to 06:00 (12th day). The collection time-points were set to include 10:00, i.e., 60 mins into the training time to detect gene expression that has been synchronized with the training time (*18, 19*). All foragers collected at 06:00 were stationary but “*active*”, 10:00, 14:00 were “*active*” (*52*), 18:00 were mostly immobile but more active than those at 06:00, 22:00 and 02:00 which were in “*second*” or “*third*” sleep stages (*52*). All collections apart from that at 10:00 (done from the unrewarded feeder plate) were done from the hive by temporarily taking out the comb frames from the hive. All night collections were done using dim red light. Collected foragers were immediately flash-frozen in liquid nitrogen and stored at -80°C until further processing.

### Cryo-sections of brain tissue for RNA-seq

Brains of flash-frozen bees were dissected using pre-chilled RNase-free cleaned dissection instruments. To avoid tissue thawing the embedding mold filled with tissue-freezing media (OCT; Leica Microsystems 3P, 14020108926) was slowly cooled on dry ice and dissected brains were placed into the mold once the lower half of the OCT was firmly frozen. Tissue block was allowed to be frozen completely on dry ice.

Before sectioning cryomolds containing brains were equilibrated to the cryostat (SLEE MEV) chamber temperature of -28°C for at least 30 minutes. Each brain was sectioned at 10μm thickness. Serial cryosections were mounted on a 45 mins UV disinfected PEN membrane slide (ThermoFisher Scientific Catalogue no. LCM0522). Within 8 mins from the mounting of the first section the slides were transferred to a slide box containing desiccant beads kept inside a dry ice container. None of the sections were allowed to dry at RT, to preserve RNA integrity.

### Tissue staining and Laser capture microdissection (LCM)

Tissue dehydration and staining was performed according to manufacturer’s protocol for (Arcturus Histogene Frozen Section Staining Kit; (KIT0401) with some alterations in timing of processing steps. LCM was performed with Zeiss PALM MicroBeam Laser Microdissection System. Marked regions of interest, ROIs (sKCs) on each section were cut and captured using system’s pulsed UV laser (337nm), with optimum energy and focus, set for the given sample. No more than 20-25 minutes were spent for each slide.

### Sample pooling, RNA isolation, RNA quality check (QC)

The sKC-enriched tissue samples from one brain captured by LCM (total of 314 brains) were collected into adhesive caps of microcentrifuge tubes (Zeiss AdhesiveCap 200 clear). Then tissues were immediately lysed and homogenized using Qiagen RNeasy Plus Micro Kit components (Catalogue no. 74034). Manufacturer’s protocol was followed and microtubes with tissue lysates were transferred to dry ice and subsequently stored at -80°C till further processing to isolate RNA. Frozen tissue lysates from 11-12 individual brains per replicate of a time-point were thawed at 37°C for 5 mins. Then, samples were pooled and further steps of RNA isolation were done according to manufacturer’s instructions. A total of 27 out of 28 samples (7 time-points x 2 pooled replicates x 2 colonies/behavior experiments) were submitted for mRNA library preparation and RNA sequencing at NCBS-TIFR Next Gen Sequencing facility. The final number of samples was 27 as one replicate was lost during sample processing due to instrumental malfunction.

### RNA-seq experimental details and data analysis

mRNA was isolated from extracted whole RNA using NEBNext® Poly(A) mRNA Magnetic Isolation Module (Catalogue no. E7490L). Stranded mRNA libraries were prepared with NEBNext® Ultra™ II Directional RNA Library Prep with Sample Purification Beads (Catalogue no. E7765L). mRNA library quality was evaluated for all samples with High Sensitivity D1000 Screen Tape ®, Agilent Technologies. Paired end RNA-seq data were obtained from Illumina Hiseq2500 platform using 2X100 bp sequencing read length. A total number of paired-end reads generated in a run was 851 and total number of million reads per sample ranged from 27.02 to 37.44.

Raw RNA-seq reads were trimmed to remove adaptors using Trimmomatic (*53*). FastQC (*54*) was done for each sample pre- and post-trimming. Reads were mapped to *Apis mellifera* genome (NCBI Assembly Amel_HAv3.1) using STAR RNA-seq aligner (*55*). The number of reads aligning to each gene was counted using featureCounts (*56*). To determine the number of detected genes, we set a filtering-criteria = minimum count value of 5 in one or more replicates per time-point, and with that we detected 9229 genes. The DESeq2 package within R (*57*) was used for differential expression analyses, identifying 995 significantly differentiated genes (DEGs) through pairwise comparisons across all time points with an adjusted p-value (p.adj) threshold of less than 0.05. These DEGs showed significant change (p.adj < 0.05) in their expression levels between at least two time points. Additionally, DESeq2 was utilized to obtain normalized count values for these selected DEGs, and subsequently, these normalized counts were scaled to generate heatmap visualizations. The MetaCycle package within R was used to evaluate circadian oscillation among the detected genes (*58*). Fisher’s exact test was used to detect the number of genes showing daily periodicity with BH.Q. values of either < 0.3 or 0.05 as correspondingly indicated. To generate heatmap visualizations for MetaCycle genes, normalized count values were scaled. The GO analysis was done using ShinyGO (*59*) with parameters as mentioned in the legends corresponding to the respective figures.

### RNA in-situ hybridization with RNAscope Technology

Brain samples for RNAscope study (an advanced type of smFISH; *60, 61*) were collected from time-trained foragers of a third honey bee colony that was exposed to the same time-training procedure as in the RNA-seq study but conducted in November 2022. On the collection day (Day 11) the feeder was rewarded to visualize target gene expression patterns in the brain during continuous food-rewarded foraging (*62*). Collected brains were serial sectioned at 12μm thickness using the same parameters as used for the LCM sectioning, and mounted on Fisherbrand; Superfrost Plus microscope slides (Catalogue no. 1255015). The smFISH with RNAscope technology using the Multiplex Fluorescent Reagent Kit v2 (Catalogue no. 323100) was performed according to manufacturer’s protocol for fresh-frozen samples from Advanced Cell Diagnostics (ACD), a Bio-Techne Brand with minor alteration in the duration spent for fixing sections with 4% paraformaldehyde. The manufacturer (ACD) designed 20 ZZ RNAscope probes target all NCBI annotated transcript variants, and the targeted nucleotide sequences for the respective genes, but only their coding regions (exons). The details of fluorophores used are summarized in the table below. Slides were coverslipped after counterstaining with H-VECTASHIELD Antifade Mounting Medium with DAPI (Vector Laboratories, Catalogue no. H-1200-10).

Fluorophores

| Specification | Catalogue No. |
| --- | --- |
| TSA CYANINE 5 SYSTEM | NEL705A001KT (50-150 SLIDES, Akoya Biosciences) |
| TSA CYANINE 3 SYSTEM | NEL704A001KT (50-150 SLIDES, Akoya Biosciences) |
| TSA Vivid Fluorophore 520 | 323271 (ACD, Bio-Techne Brand) |

### Confocal microscopy and image analysis

All samples were imaged using an Olympus FluoView 3000 confocal microscope. All imaging parameters like dwell time, laser power, and gain were kept constant across samples. Representative confocal images were adjusted for brightness and contrast (evenly across images) in Fiji software (ImageJ, *63*). All quantifications were done in Fiji. Area analysis was done by selecting the ROI, in only DAPI channel and then the same ROI was projected to all the other channels. For area analysis the raw image was median filtered and then thresholded with dark background following which area and area fraction were measured. The plots were generated in GraphPad Prism 10. In GraphPad Prism 10 the data were checked for normality (alpha=0.05), following which one-way ANOVA (test for linear trend) was used to determine whether the means for each time-point increases gradually from one time-point to the others, P value < 0.05. For quantification of correlation, JACoP plugin was used and Pearson’s coefficient (r), and Manders’ coefficients (M1 and M2) were determined from raw images.

