## Supplementary figures and table for "Multiple molecular links between the circadian clock and memory centers in honey bees"

Supplementary materials

**A** Network for all DGs (n = 9229)

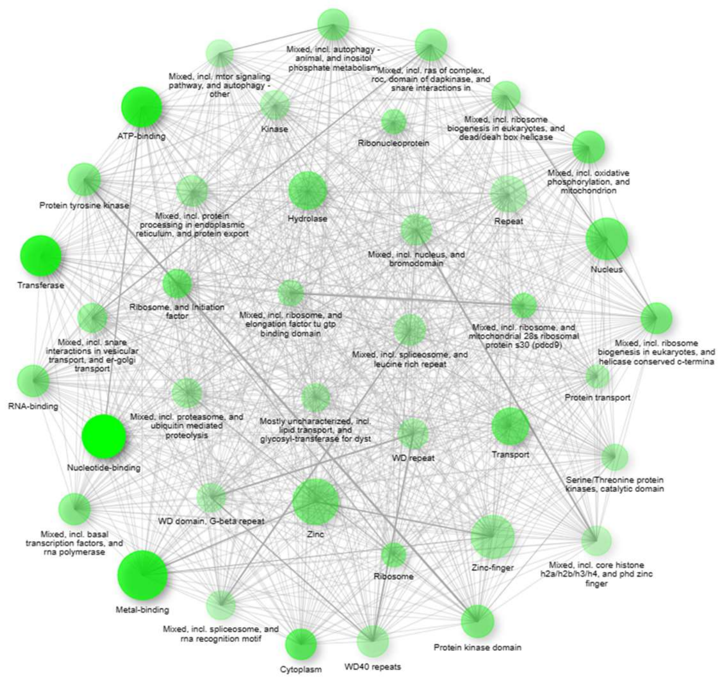


**B** Bar plot for all DGs (n = 9229)

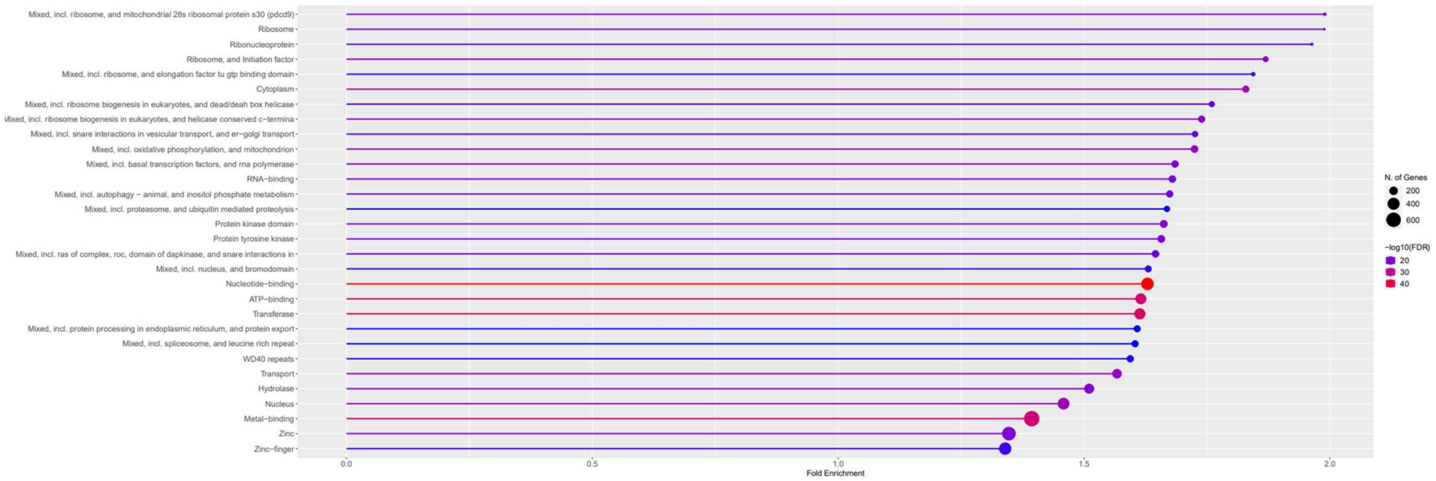


**Fig. S1. Gene interaction analysis of detected gene (DGs) sets.**

(A) Illustration of gene networks for all detected genes showing relation between enriched pathways.

(B) Bar plot with all detected genes showing no. of genes in each category with FDR < 0.01.

All plots are predicted by GO analysis. For gene network, two nodes (pathways) are connected when 20% or more genes are shared. Darker nodes are more significantly enriched gene sets. Bigger nodes represent larger gene sets. Thicker edges represent more overlapped genes.

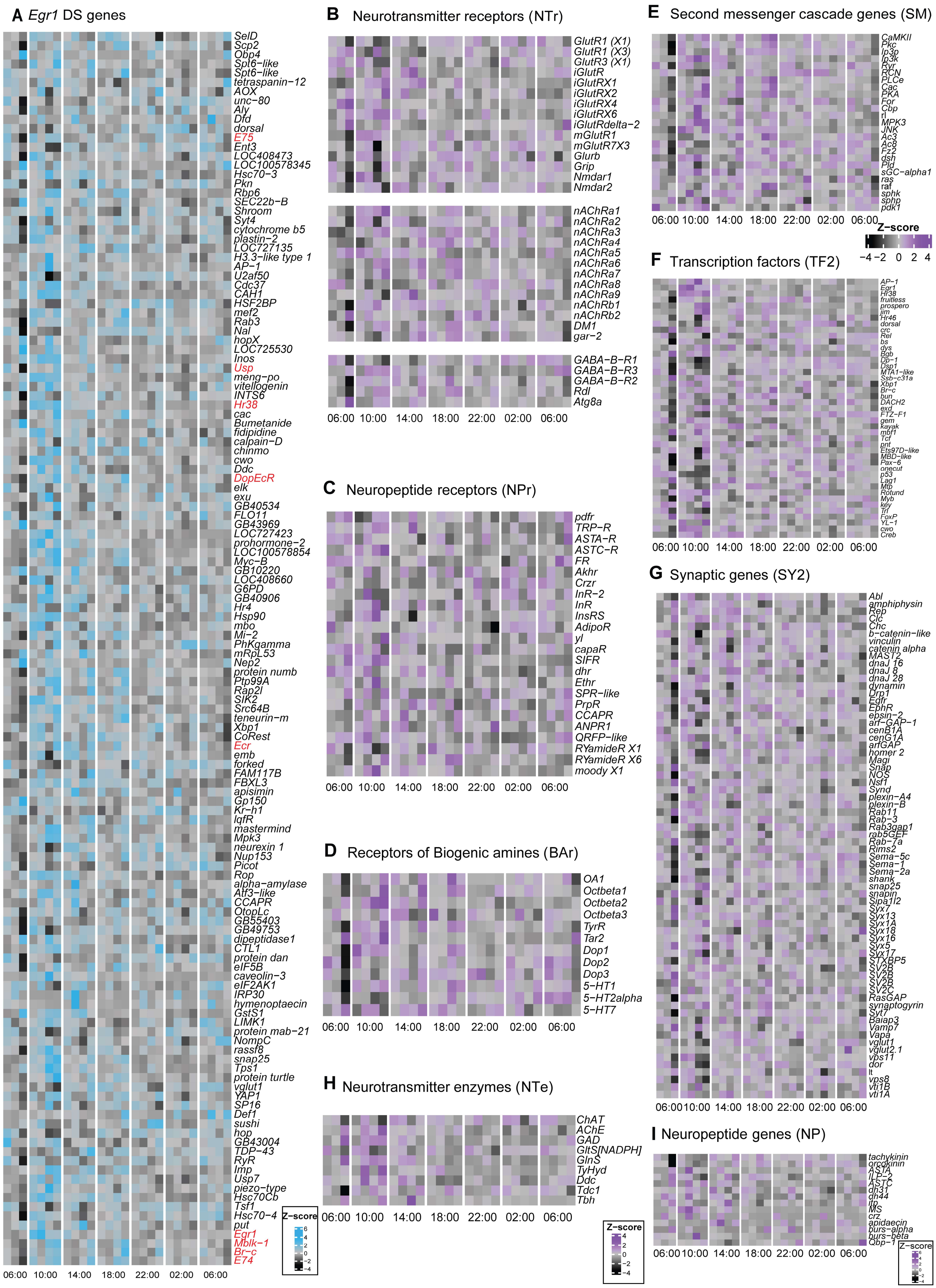
A Egr1 DS genes
SelD
Scp2
Obp4
Spt6−like
Spt6−like
tetraspanin−12
AOX
unc−80
Aly
Dfd
dorsal
E75
Ent3
LOC408473
LOC100578345
Hsc70−3
Pkn
Rbp6
SEC22b−B
Shroom
Syt4
cytochrome b5
plastin−2
LOC727135
H3.3−like type 1
AP−1
U2af50
Cdc37
CAH1
HSF2BP
mef2
Rab3
Nal
hopX
LOC725530
Inos
Usp
meng−po
vitellogenin
INTS6
Hr38
cac
Bumetanide
fidipidine
calpain−D
chinmo
cwo
Ddc
DopEcR
elk
exu
GB40534
FLO11
GB43969
LOC727423
prohormone−2
LOC100578854
Myc−B
GB10220
LOC408660
G6PD
GB40906
Hr4
Hsp90
mbo
Mi−2
PhKgamma
mRpL53
Nep2
protein numb
Ptp99A
Rap2l
SIK2
Src64B
teneurin−m
Xbp1
CoRest
Ecr
emb
forked
FAM117B
FBXL3
apisimin
Gp150
Kr−h1
IqfR
mastermind
Mpk3
neurexin 1
Nup153
Picot
Rop
alpha−amylase
Atf3−like
CCAPR
OtopLc
GB55403
GB49753
dipeptidase1
CTL1
protein dan
eIF5B
caveolin−3
eIF2AK1
IRP30
hymenoptaecin
GstS1
LIMK1
protein mab−21
NompC
rassf8
snap25
Tps1
protein turtle
vglut1
YAP1
SP16
Def1
sushi
hop
GB43004
TDP−43
RyR
Imp
Usp7
piezo−type
Hsc70Cb
Tsf1
Hsc70−4
put
Egr1
Mblk−1
Br−c
E74
06:00 10:00 14:00 18:00 22:00 02:00 06:00
Z−score
6
4
2
0
−2
−4
B Neurotransmitter receptors (NTr)
GlutR1 (X1)
GlutR1 (X3)
GlutR3 (X1)
iGlutR
iGlutRX1
iGlutRX2
iGlutRX4
iGlutRX6
iGlutRdelta−2
mGlutR1
mGlutR7X3
Glurb
Grip
Nmdar1
Nmdar2
nAChRa1
nAChRa2
nAChRa3
nAChRa4
nAChRa5
nAChRa6
nAChRa7
nAChRa8
nAChRa9
nAChRb1
nAChRb2
DM1
gar−2
GABA−B−R1
GABA−B−R3
GABA−B−R2
Rdl
Atg8a
06:00 10:00 14:00 18:00 22:00 02:00 06:00
C Neuropeptide receptors (NPr)
pdfr
TRP−R
ASTA−R
ASTC−R
FR
Akhr
Crzr
InR−2
InR
InsRS
AdipoR
yl
capaR
SIFR
dhr
Ethr
SPR−like
PrpR
CCAPR
ANPR1
QRFP−like
RYamideR X1
RYamideR X6
moody X1
06:00 10:00 14:00 18:00 22:00 02:00 06:00
D Receptors of Biogenic amines (BAr)
OA1
Octbeta1
Octbeta2
Octbeta3
TyrR
Tar2
Dop1
Dop2
Dop3
5−HT1
5−HT2alpha
5−HT7
06:00 10:00 14:00 18:00 22:00 02:00 06:00
H Neurotransmitter enzymes (NTe)
ChAT
AChE
GAD
GltS[NADPH]
GlnS
TyHyd
Ddc
Tdc1
Tbh
06:00 10:00 14:00 18:00 22:00 02:00 06:00
Z−score
4
2
0
−2
−4
E Second messenger cascade genes (SM)
CaMKII
Pkc
Ip3p
Ip3k
Ryr
RCN
PLCe
Cac
PKA
For
Cbp
rl
MPK3
JNK
Ac3
Ac8
Fz2
dsh
Pld
sGC−alpha1
ras
raf
sphk
sphp
pdk1
06:00 10:00 14:00 18:00 22:00 02:00 06:00
Z−score
−4 −2 0 2 4
F Transcription factors (TF2)
AP−1
Egr1
Hr38
fruitless
prospero
jim
Hr46
dorsal
crc
Rel
bs
dys
Bgb
DP−1
Dsp1
MTA1−like
Ssb−c31a
Xbp1
Br−c
bun
DACH2
exd
FTZ−F1
gem
kayak
mbf1
Tcf
pnt
Ets97D−like
MBD−like
Pax−6
onecut
p53
Lag1
Mtp
Rotund
Myb
key
Trl
FoxP
YL−1
cwo
Creb
06:00 10:00 14:00 18:00 22:00 02:00 06:00
G Synaptic genes (SY2)
Abl
amphiphysin
Rep
Clc
Chc
b−catenin−like
vinculin
catenin alpha
MAST2
dnaJ 16
dnaJ 8
dnaJ 28
dynamin
Drp1
Egfr
EphR
epsin−2
arf−GAP−1
cenB1A
cenG1A
arfGAP
homer 2
Magi
Snap
NOS
Nsf1
Synd
plexin−A4
plexin−B
Rab11
Rab−3
Rab3gap1
rab5GEF
Rab−7a
Rims2
Sema−5c
Sema−1
Sema−2a
shank
snap25
snapin
Sipa1l2
Syx7
Syx13
Syx1A
Syx18
Syx16
Syx5
Syx17
STXBP5
SV2B
SV2B
SV2B
SV2C
RasGAP
synaptogyrin
Syt7
Baiap3
Vamp7
Vapa
vglut1
vglut2.1
vps11
dor
lt
vps8
vti1B
vti1A
06:00 10:00 14:00 18:00 22:00 02:00 06:00
I Neuropeptide genes (NP)
tachykinin
orcokinin
ASTA
ILP−2
ASTC
dh31
dh44
itp
MS
crz
apidaecin
burs−alpha
burs−beta
Obp−1
06:00 10:00 14:00 18:00 22:00 02:00 06:00
Z−score
6
4
2
0
−2
−4

**Fig. S2. Expression pattern of categories of *Egr1* DS- and NS- library genes.**

(A) *Egr1* DS genes. Ecdysteroid signaling genes are marked in red.

(B) Receptors of neurotransmitters (NTr).

(C) Receptors of neuropeptides (NPr).

(D) Receptors of biogenic amines (BAr).

(E) Second messenger cascade genes (SM).

(F) Transcription factors (TF2).

(G) Synaptic genes (SY2).

(H) Neurotransmitter enzymes (NTe).

(I) Neuropeptide genes (NP).

**A** Network for all DEGs (n = 995)

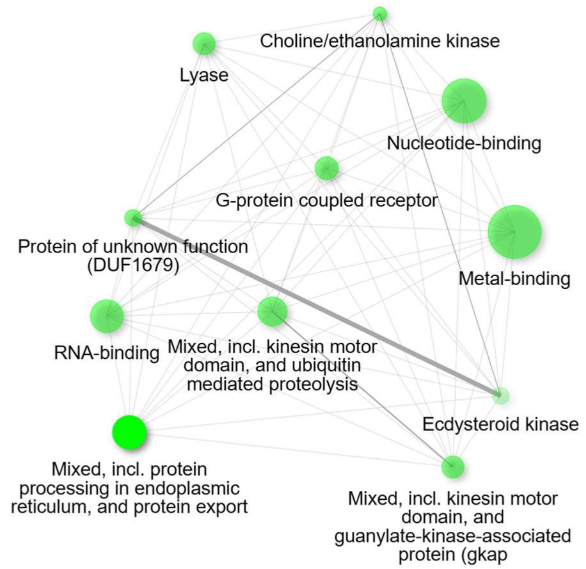


**B** Bar plot for all DEGs (n = 995)

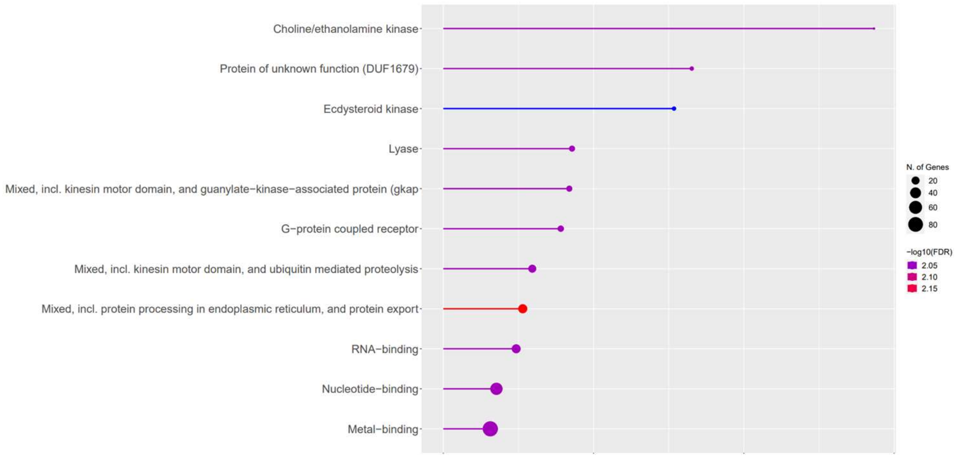


**Fig. S3. Gene interaction analysis of differentially expressed gene sets.**

(A) Illustration of gene networks for all DEGs.

(B) Bar plot for DEGs showing no. of genes assigned to each category.

All plots are predicted by GO analysis. For gene network, two nodes (pathways) are connected when 20% or more genes are shared. Darker nodes are more significantly enriched gene sets. Bigger nodes represent larger gene sets. Thicker edges represent more overlapped genes.

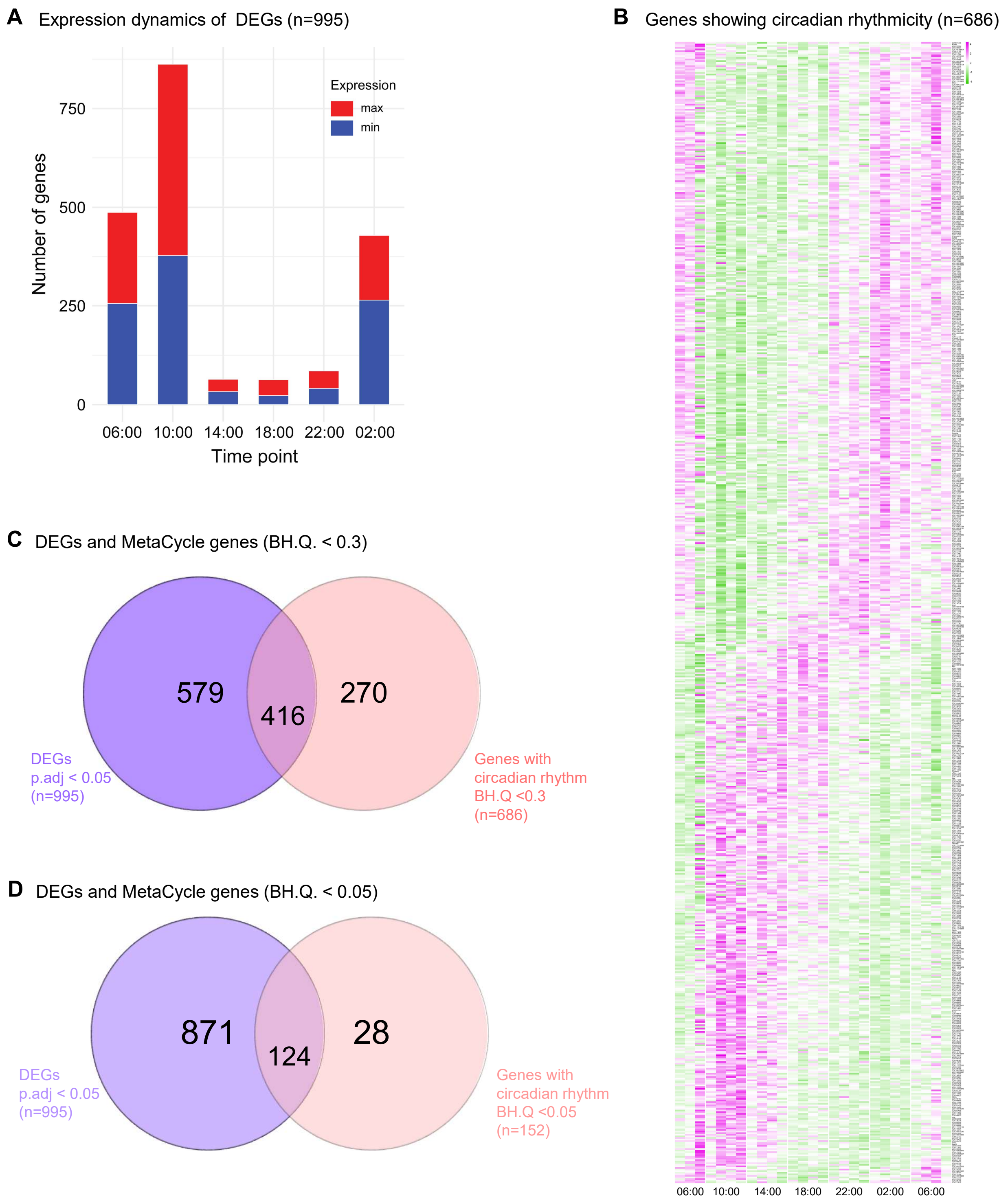


**Fig. S4. Dynamics of gene expression in sKC-enriched RNA-seq dataset.**

(A) Bar chart for 995 DEGs showing no. of genes (y-axis) expressed at their max (red) and min (blue) in each time-point (x-axis). The genes for day 11, 06:00 and day 12, 06:00 are combined as no genes were found to be differentially expressed between these two time points (pairwise comparison test, p.adj <0.05). Three peaks are observed, main peak at 10:00; one hour into the training-time (feeder not rewarded), smaller peak at 06:00 (hits of lights on), another peak at 02:00 (late sleep stages).

(B) Heatmap showing gene expression dynamics of the ones having circadian rhythmicity. MetaCycle algorithm, Fisher's exact test, BH.Q < 0.3

(C) Venn diagram showing the overlap among the genes showing differential expression (p.adj< 0.05) and genes showing circadian oscillation (BH.Q. < 0.3).

(D) Venn diagram showing the overlap among the genes showing differential expression (p.adj< 0.05) and genes showing circadian oscillation (BH.Q. < 0.05).

**A** Clock-related neuropeptides

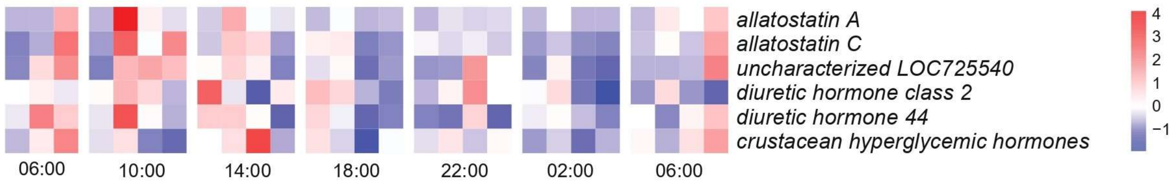


**Fig. S5. Dynamics of clock-related neuropeptide genes.**
(A) Heatmaps showing expression patterns of clock-related neuropeptide genes [predicted in (*35*) and detected in our RNA-seq dataset.

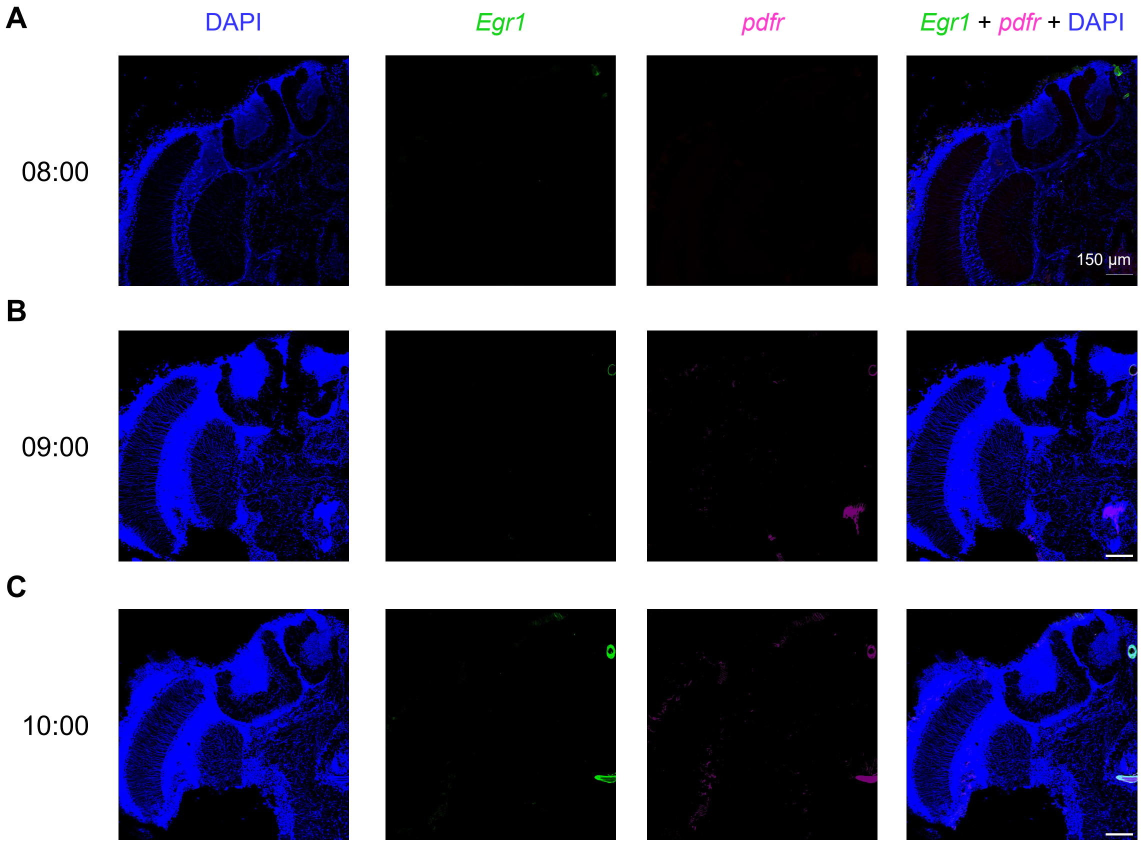


**Fig. S6. Specificity of ISH with RNAscope and that of both *Egr1* and *pdfr* probes validated with no staining in negative controls.**
(A, B and C) Respective negative control images for 08:00, 09:00 and 10:00. All reagents for the ISH with RNAscope were same except for the antisense probes specific to the genes of interest. This confirms the specificity of the technique and the probes. Scale bars, 150 µm. Time-points 08:00 or 09:00, and 10:00 as indicated.

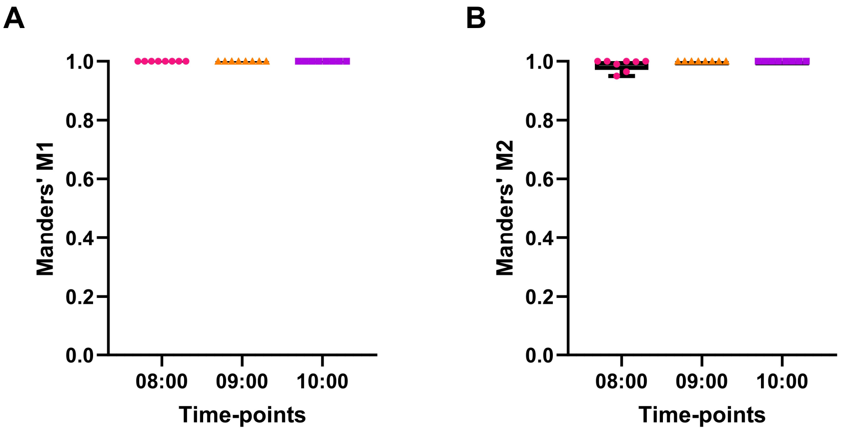


**Fig. S7. Co-expression of *pdfr* and *Egr1* in KCs of time-trained foragers at 08:00, 09:00 and 10:00.** (A) Box-whiskers plot showing Manders' correlation coefficient (M1) suggesting high fraction (M1=1) of expression of *pdfr* in *Egr1* positive cells. (B) Box-whiskers plot showing Manders' correlation coefficient (M2) suggesting high fraction (M2=1) of expression of *Egr1* in *pdfr* positive cells. Quantified for N=3, different calyces at high magnification, brain section depth: Mid, 150 to 300 μm.

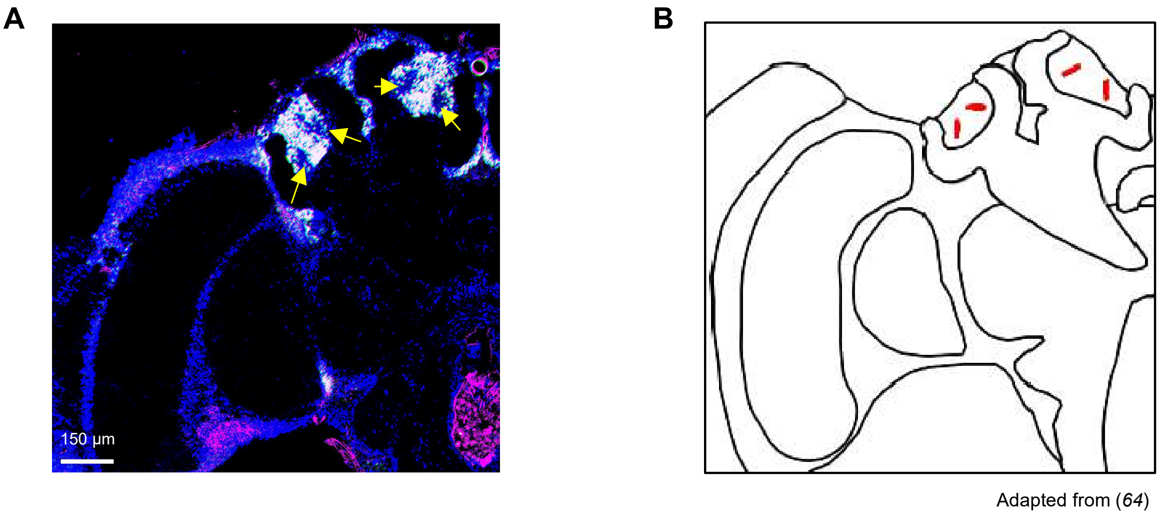


**Fig. S8. Lack of co-expression of *Egr1* and *pdfr* in *foxP*⁺ region unlike most KCs of time-trained foragers at 10:00.**
(A) Absence of *Egr1* and *pdfr* co-expression from *foxP*⁺ region (yellow arrows) even when most KCs show co-expression at 10:00 during continuous foraging. Scale bar, 150 µm. (B) Schematic representation of *foxP*⁺ region as identified in (*64*) and marked in red. Brain section depth: Mid, 150 to 300 µm.

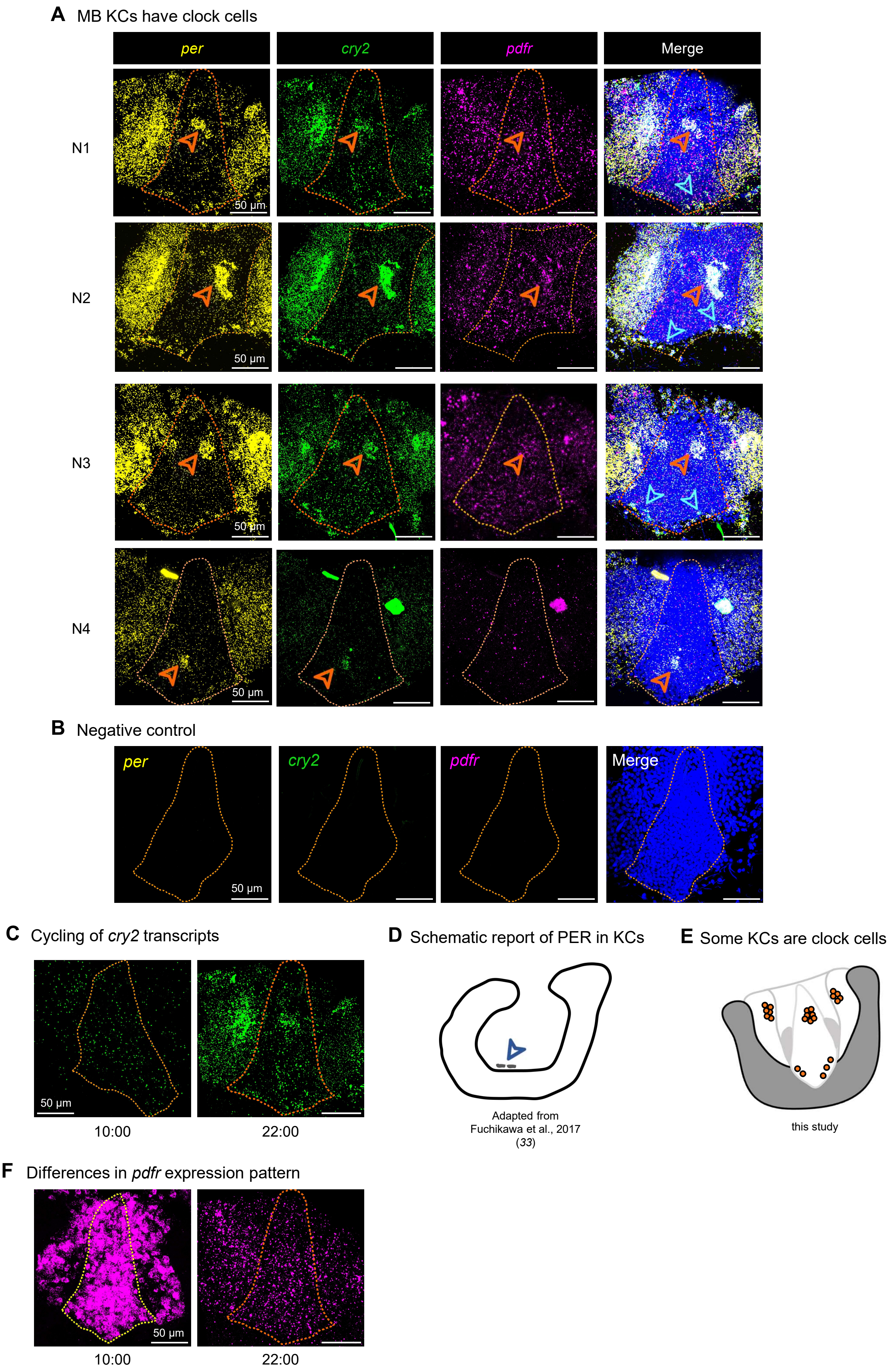
A MB KCs have clock cells
per
cry2
pdfr
Merge
N1
N2
N3
N4
50 µm
B Negative control
per
cry2
pdfr
Merge
50 µm
C Cycling of cry2 transcripts
50 µm
10:00
22:00
D Schematic report of PER in KCs
Adapted from
Fuchikawa et al., 2017
(33)
E Some KCs are clock cells
this study
F Differences in pdfr expression pattern
50 µm
10:00
22:00

**Fig. S9. Local circadian clock in MB KCs.**
(A) RNAscope images for *per* (yellow), *cry2* (green), and *pdfr* (magenta) and co-expression (white), when merged with DAPI (blue). sKC region, orange dotted line. Arrowheads: orange, spatial clustering of cells showing high transcript abundance; cyan, *per* (yellow) and *cry2* (green) co-expression in basal sKCs indicated earlier with immunohistochemistry with PER in (*33*). Top: One calyx of individual 1 (N1), also in Fig. 4C. Next bottom: One calyx of individual 2 (N2). Next bottom: One calyx of individual 3 (N3). Next bottom: One calyx of individual 4 (N4). Some artifacts can be found in N4, possibly due to gland (top-left) and dust particle (top-right). Both artifacts are outside the sKC region (marked with orange dotted line). The cluster in the sKCs comprise of ~560 cells across individuals
(B) Negative control images for *per* (yellow), *cry2* (green), and *pdfr* (magenta). All reagents for the ISH with RNAscope were same except for the antisense probes specific to the genes of interest. This confirms the specificity of the technique and the probes. (C) RNAscope images for *cry2* expression at 10:00 (left) and 22:00 (right), the highest and the lowest expression time-points (predicted by RNA-seq). The observed expression pattern confirms cycling of *cry2* transcripts consistent with RNA-seq results. Scale bars, 50 μm. (D) Schematic for expression of PER in basal sKCs indicated earlier with immunohistochemistry, adapted from (*33*). (E) Schematic showing clock cells in MB KCs, as interpreted from RNAscope study (cells with high transcript abundance and showing co-expression are marked in orange). (F) RNAscope images for *pdfr* expression at 10:00 (left) and 22:00 (right) indicating high and low expressions respectively.

**Table S1**: Comparison of areas showing PER immunostaining (*Fuchikawa et al. 2017*) and RNAscope with *per* probes, this study

| **Anatomical location** | **PER-ir cells (*Fuchikawa et al., 2017*, *33*; Immunostaining)** | ***per* probes (this study, RNAscope staining)** |
|---|---|---|
| $LN_1$ | yes | yes |
| $LN_2$ | yes | yes |
| DLN | yes | yes |
| DN | yes | yes |
| LO/ME | yes | yes |
| ME/LA | yes | yes |
| RE | yes | yes |
| PI | yes | yes |
| AL | yes | yes |
| MB KCs | not identified (reported as few number of PER-ir nuclei above the calyces) | yes |
| Central complex | around the central complex | neurons in close vicinity to central complex |
| SEG | no | no |
| ocelli | yes | not tested |
